# A robust and interpretable, end-to-end deep learning model for cytometry data

**DOI:** 10.1101/2020.02.05.934521

**Authors:** Zicheng Hu, Alice Tang, Jaiveer Singh, Sanchita Bhattacharya, Atul J. Butte

**Author notes:** Correspondence should be addressed to Z.H. and A.J.B.

## Abstract

Cytometry technologies are essential tools for immunology research, providing high-throughput measurements of the immune cells at the single-cell level. Traditional approaches in interpreting and using cytometry measurements include manual or automated gating to identify cell subsets from the cytometry data, providing highly intuitive results but may lead to significant information loss, in that additional details in measured or correlated cell signals might be missed. In this study, we propose and test a deep convolutional neural network for analyzing cytometry data in an end-to-end fashion, allowing a direct association between raw cytometry data and the clinical outcome of interest. Using nine large CyTOF studies from the open-access ImmPort database, we demonstrated that the deep convolutional neural network model can accurately diagnose the latent cytomegalovirus (CMV) in healthy individuals, even when using highly heterogeneous data from different studies. In addition, we developed a permutation-based method for interpreting the deep convolutional neural network model and identified a CD27-CD94+ CD8+ T cell population significantly associated with latent CMV infection. Finally, we provide a tutorial for creating, training and interpreting the tailored deep learning model for cytometry data using Keras and TensorFlow (github.com/hzc363/DeepLearningCyTOF).

## Main

Modern cytometry technologies, including flow cytometry and mass cytometry (CyTOF), are able to characterize cell mixtures at the single-cell resolution with over 40 markers^1^. Multi-dimensional cytometry data contains rich information that can be used to identify key cellular changes induced by diseases or other perturbations, such as viral infections, cancer immunotherapies, and vaccinations^2–4^. In addition, cytometry measurements have been utilized for decades to diagnose a variety of conditions, such as leukemia, allergies, and infectious diseases^5–7^.

The analysis of cytometry data typically starts with identifying cell populations by manual gating or by automated clustering using computational methods, including FLOCK, MetaCyto, flowSOM, and others^8–10^. The subsequent analysis then uses summary statistics of the identified cell populations, including abundance and mean marker expression levels, to identify disease-associated cells or to predict clinical outcomes^11,12^. This approach is an intuitive way to analyze cytometry data and has yielded highly interpretable results. However, the approach has several disadvantages. First, in the cell gating step, the original cytometry data is reduced to summary statistics of cell subsets, potentially leading to the loss of important information such as the correlation between cell markers and the distribution of marker expression within each cell subset. Second, the commonly used approach requires all samples to be clustered in the same way, making it sensitive to batch effects and the choice of clustering methods. Finally, the approach may fail to detect cellular changes that do not lead to distinct cell populations, such as the continuous up-regulation of CTLA-4 in T cells in response to varying degrees of stimulation^13^.

Several recent studies have explored alternative approaches to analyze cytometry data, bypassing the requirement for cell gating or cell clustering. We previously developed CytoDx, which fits the cytometry data using a two-stage linear model^14^. Another study developed CellCNN to model the cytometry data using convolutional neural networks^15^. Both of these methods utilize the full cytometry data, rather than the summary statistics from cell gating steps, therefore are more advantageous for disease diagnosis and identification of disease-associated cells^14,15^. On the other hand, these existing methods still use relatively simple models (linear regression and neural networks with a single convolutional layer). Both are only capable of combining cell markers linearly at the single-cell level, thus preventing them from capturing more complex combinatorial cellular phenotypes in cytometry measurement data.

The interpretation of the CytoDx and CellCNN models also remained a challenge. The methods developed in previous studies can only interpret parts of the models. To identify cell populations that are associated with outcomes of interest, both methods leverage the one-to-one correspondence between cells and the intermediate output of the model (the output of the cell-level model in CytoDx and convolutional layers in CellCNN). New methods are required to extract biological insights from the full models.

In this study, we developed and tested a framework for modeling cytometry data using a deep convolutional neural network (CNN), in which multiple hidden layers are used to model the high-dimensional cytometry data. Leveraging multiple large publicly-available CyTOF datasets (472 samples from 9 studies) available in ImmPort^16–19^, we demonstrate that the deep CNN model is able to diagnose asymptomatic cytomegalovirus infection with high accuracy, even in the presence of strong heterogeneity between datasets. In addition, we developed a permutation-based method to interpret the full deep CNN model. We identified a previously undescribed CD27− CD94+ CD8+ T cell population that is significantly increased in subjects with latent CMV infections, across nine studies. Interestingly, the CD27− CD94+ subset is increased in all four compartments (Naïve, effector, effector memory and central memory) of CD8+ T cells, suggesting that CMV infection induces the CD94+ CD27− phenotype through a mechanism that is distinct from T cell activation and memory.

## Results

### A deep convolutional neural network for cytometry data

We designed a deep convolutional neural network architecture tailored to the cytometry data. The input into the model is the raw cytometry data, which are matrices with rows representing cells and columns representing markers. The outputs of the model are sample-level information of interest, such as disease diagnosis, drug responsiveness or the presence of a genetic deficiency. The internal layers of the deep CNN model include multiple convolutional layers to extract cell-level features, a pooling layer to aggregate the cell-level features into sample level features, and dense layers to capture the interaction between the sample-level features (**Fig. 1**).

**Figure 1:**
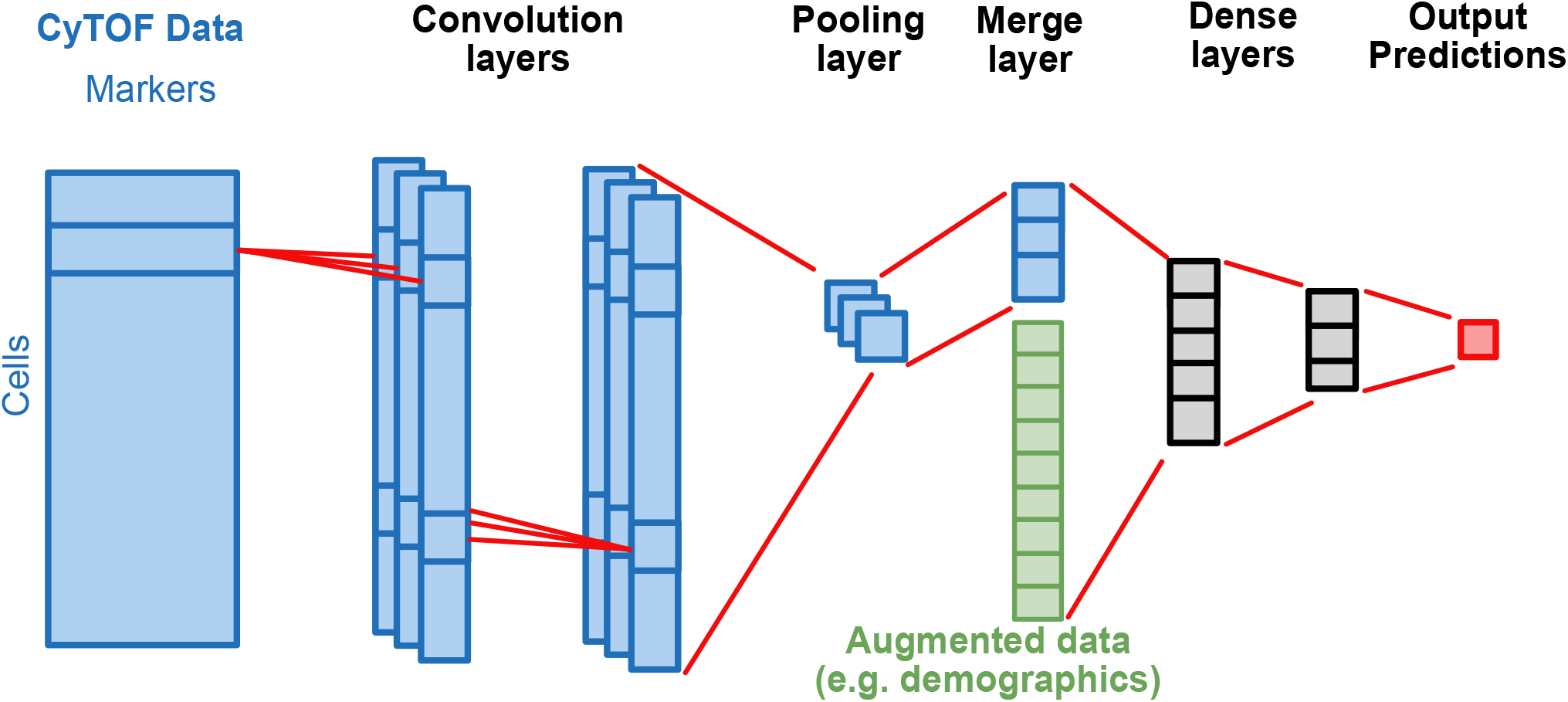
Schematic diagram showing the structure of the deep CNN model. The model takes arcsinh-transformed cytometry data (dimension equals to Num. of cells × Num. of Markers) as input, extract cellular features using convolution layers (filter size equals to 1 × Num. of Markers) and aggregates cellular features using max or average pooling. The aggregated features can be augmented with other non-cytometry data. The dense layers combine the augmented data and predict the outcome of interest, which can be either continuous or categorical variables.

A key characteristic of cytometry data is that it represents an unordered collection of cells. The data representation is similar to the point cloud in computer vision^20^. In order to model this type of data in an efficient way, the neural network needs to be invariant to the permutation of rows in the data^21^. We achieved this by 1) designing “one-cell” filters in convolutional layers, which combines all marker information within the same row, but not across rows. 2) applying either max or mean function over all cells in the pooling layer, both of which are invariant to the permutation of data.

In addition to cytometry data, the deep CNN model allows the incorporation of external information, such as demographics (age, gender, and race) and results from other experiments. Specifically, the output of the pooling layer can be combined with other sample-level information to improve model performance and to adjust for control variables (**Fig. 1**).

### The deep CNN model accurately predicts asymptomatic CMV infection

To test the performance of the deep CNN model, we applied it to nine CyTOF datasets to train it to diagnose asymptomatic cytomegalovirus (CMV) infection. The dataset spans nine human immunology studies and contains 596 peripheral blood mononuclear cells (PBMC) samples from 313 subjects^16–19^. We split the nine studies into training, validation, and testing datasets. To ensure an unbiased performance evaluation, we selected SDY515 and SDY519 as validation and testing datasets, which do not share subjects with other studies (**Fig. 2**).

**Figure 2:**
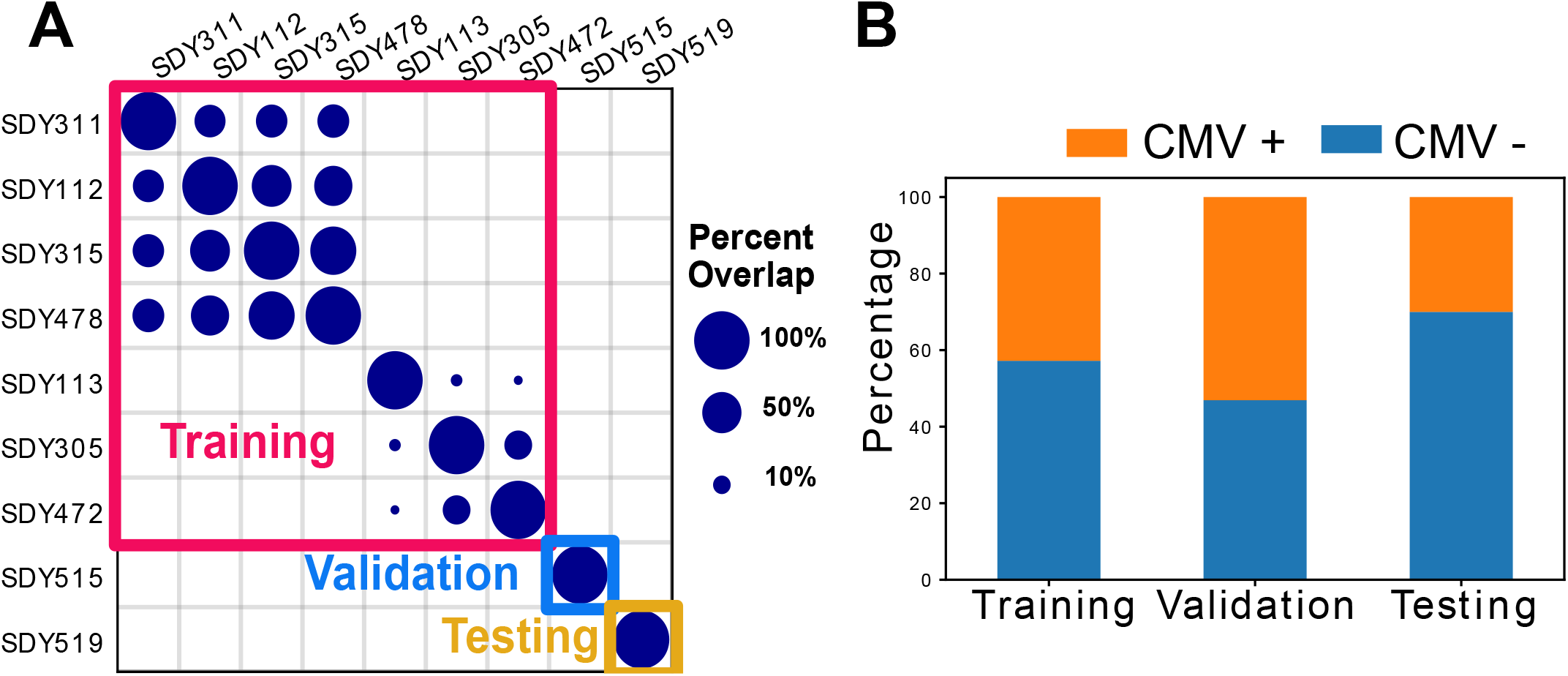
Overview of the CyTOF dataset. (**A**) Overlap of subjects between nine studies and the split of the studies into training, validation and testing datasets. The dot size represents the percentage of overlapping subjects between studies. (**B**) percentages of CMV positive and CMV negative individuals in training, validation and testing datasets.

We trained and optimized the deep CNN model using training and validation datasets. The final model is evaluated using the test dataset. The deep CNN model is able to diagnose the CMV infection with high accuracy (Area Under the Receiver Operating Curve (AUROC) = 0.94, Area Under the Precision-Recall Curve (AUPRC) = 0.91). To benchmark the performance of the deep CNN model, we trained and tested several existing methods, including CytoDx, CellCNN and FlowSOM^10,14,15^. The 1000-fold Bootstrap analysis shows that the deep CNN model outperforms the existing methods (**Fig. 3**).

**Figure 3:**
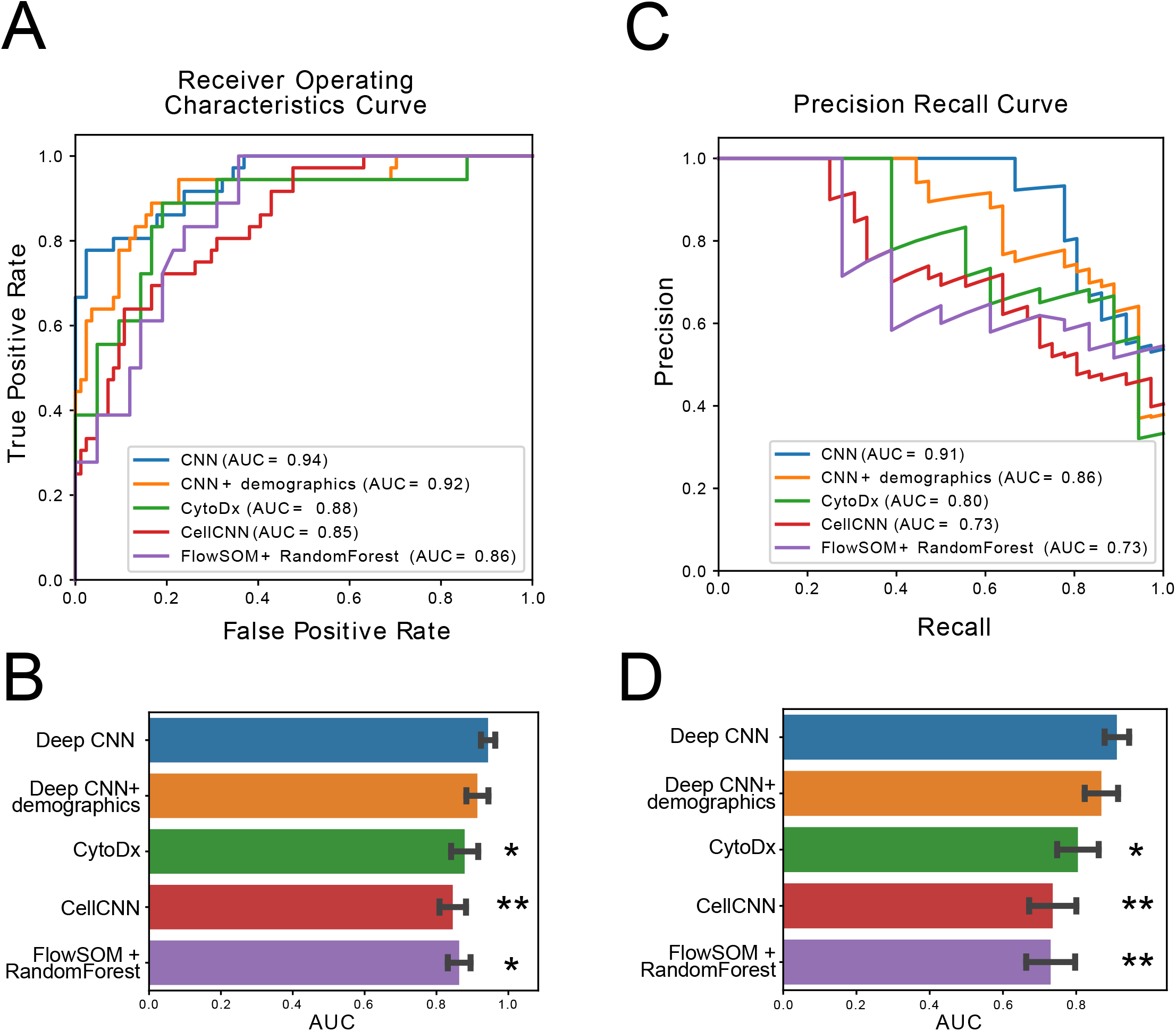
The performance of deep CNN and other methods. We used Deep CNN, CytoDx, CellCNN and FlowSOM to diagnose latent CMV in the test dataset. (**A**) The performances of the models measured by the Receiver-Operator Characteristics curves. (**B**) The areas under the receiver-operator characteristics curves. The error bars represent the standard deviation. The standard deviation and p values are measured by 1000 fold bootstrapping. (**C**) The performances of the models measured by the Precision-Recall Curves. (**D**) The areas under the Precision-Recall Curves. The error bars represent the standard deviation. The standard deviation and p values are measured by 1000 fold bootstrapping. *, p value < 0.05; **, p value < 0.01.

We tested the robustness of the model against the choice of training, validation and testing dataset. In each iteration, we randomly assigned one study as the validation dataset, one study as the testing dataset and the rest studies as the training dataset to train and evaluate the deep CNN model. We repeated the process 10 times and found that the model is able to diagnose CMV accurately in all iterations (AUROC ranges from 0.93 to 0.97, see **Table S1**).

Previous studies have demonstrated that the CMV prevalence is significantly different between age, sex, and race groups^22–24^. Therefore, augmenting the CyTOF data with demographic data can potentially improve the performance of the deep CNN model. We tested the augmented model and found that its performance is similar to the non-augmented model, suggesting that demographics data does not provide additional information to the model in this particular case (**Fig. 3**).

### The deep CNN model mitigates batch effects across studies

Visual inspection reveals an obvious heterogeneity between CyTOF data from different studies (**Fig. 4A**), which is likely due to a combination of technical differences during data acquisition and the biological differences between study cohorts. Despite the heterogeneity, the deep CNN model is able to accurately diagnose CMV infection in all nine datasets, suggesting that the model is able to extract CMV related signals from noises caused by batch effects and other non-CMV related differences in the immune system. We measured the cross-study heterogeneity using a Kruskal-Wallis test in each layer of the deep CNN model. We found that the heterogeneity is gradually mitigated across the layers of the deep CNN model (**Fig. 4B-G)**. The heterogeneity is the strongest at the input layer (p-value = 4.7 × 10^−74^) but became insignificant in the output layer (p-value = 0.16). Notably, the heterogeneity is not only reduced among the studies within the training dataset but also mitigated across the training, validation and test dataset. The results suggest that the deep CNN model is robust and is generalizable to data outside the training dataset.

**Figure 4:**
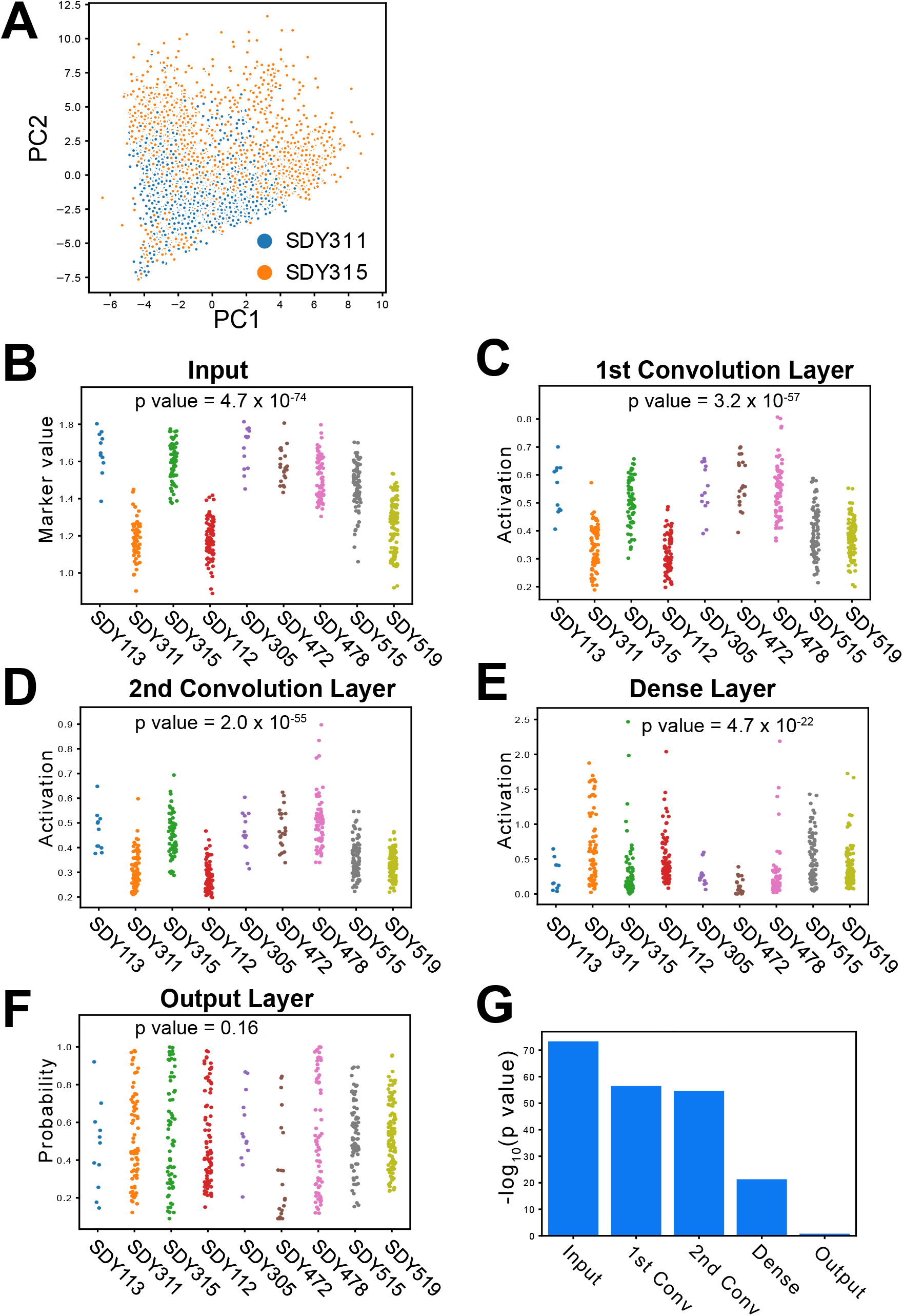
The deep CNN model mitigates the heterogeneity in cytometry data. (**A**) A PCA plot visualizing the batch effect between cytometry data from SDY311 and SDY315. (**B-F**) Dot plots showing the mean values in different layers of the deep CNN model, including input (**B**), first convolutional layer (**C**), second convolutional layer (**D**), pooling layer (**E**) and output layer (**F**). Each dot represents a sample in the studies. (**G**) Bar plot showing the batch effects in different layers in the deep CNN model, measured by the negative logarithm of p-values. p-values reported in **B-G** are from Kruskal–Wallis tests.

### The deep CNN model identifies novel associations between immune cell subsets and CMV infection

Leveraging the one-to-one correspondence between cells and internal nodes in the convolution layers, we first use the activation values of the convolution layers to identify cells associated with CMV infection. Using the cell definitions from the Human Immunology Project Consortium^25^, we identified 24 well-characterized cell populations from the CyTOF data. For each cell population, we quantified the mean activation value in the convolution layers. In the first convolution layer, Memory B cells, CD8+ T− Effector Memory (T-EM) cells and CD4+ T− Central Memory (T-CM) cells have the highest mean activation value from the three filters, respectively. In the second convolution layer, Effector CD8+ T cells, Plasmablasts, and CD8+ T-EM cells have the highest mean activation value from the three filters, respectively. The natural killer T (NKT) cells are also highly activated in the first filter of the second convolution layer (**Fig. 5A**). To test if the highly activated cells are associated with CMV infection, we quantified their percentage within PBMC from CMV positive and negative subjects from all nine studies. We found that two of the three cell populations (Memory B cells and CD4+ T-CM cells) activated by the first convolution layer are associated with CMV infection. The cell subsets activated in the second convolutional layers (Effector CD8+ T cells, Plasmablasts, and CD8+ T-EM cells) are all significantly associated with CMV infection (**Fig. S1**).

**Figure 5:**
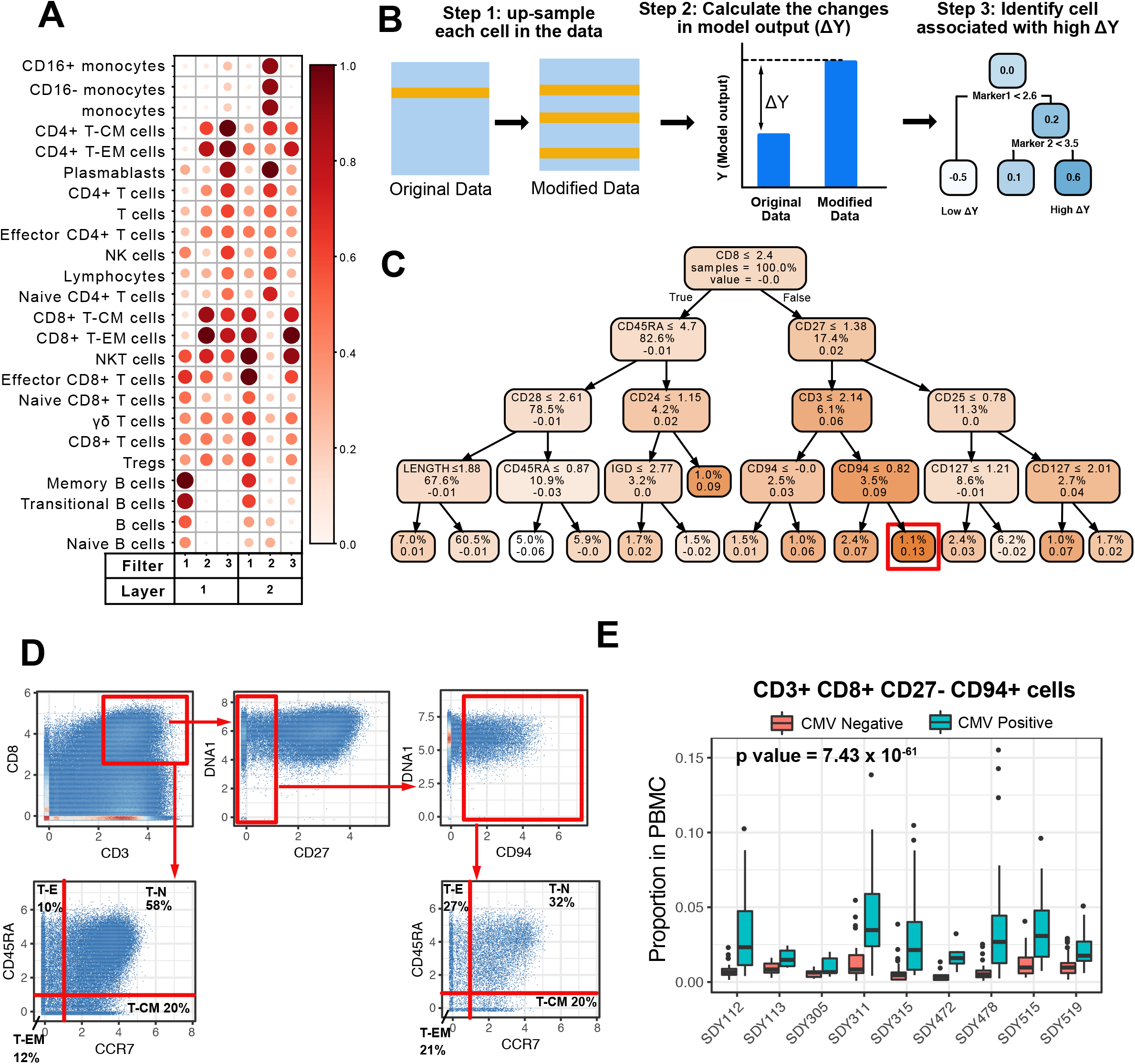
The deep CNN model identifies novel associations between immune cell subsets and CMV infection. (**A**) The mean activation value in the convolutional layers in each cell population. The activation values are normalized by dividing the activation values by the highest value in the filter. (**B**) The workflow for interpreting the full deep CNN model. (**C**) A decision tree identifies the cells that lead to the largest changes in model output (ΔY) when up-sampled. Each node represents a cell subset. The rules by which the populations split are indicated inside the nodes. The values in each node represent the percent of subset in the total population and the average change of model output (ΔY) when cells are up-sampled. The red box highlights the node with the highest mean ΔY. (**D**) Scatter plots showing the gating of the CD8+ CD3+ CD27-CD94+ cells and the composition of Naive (T-N), Effector(T-E), Effector memory (T-EM) and Central memory (T-CM) compartment in Bulk CD8+ T cells and in CD8+ CD3+ CD27-CD94+ T cells. (**E**) the percentage of CD8+ CD3+ CD27-CD94+ cells in CMV+ and CMV-subjects across nine studies. P values are from two-way ANOVA models, with CMV infection and study as two factors. The p values of the CMV infection variable are reported.

Next, we inspected beyond the convolutional layers and hope to identify the key immune differences by interpreting the full deep CNN model. We developed a permutation-based interpretation procedure (**Methods**), which is inspired by the Local interpretable model-agnostic explanations (LIME) approach^26^. Briefly, we iteratively up-sampled each cell by copying it to replace other randomly chosen cells within the sample. We then applied the deep CNN model on both the original data and the permuted data. The difference in the model output (ΔΥ) quantifies the impact of each cell on the output of the deep learning model. We then built a decision tree to identify cell subsets that have a high impact on the deep CNN model (**Fig. 5B**). We choose to use decision tree models because of their high interpretability and the structural similarity between decision trees and the hieratical cell gating.

The decision tree identifies a CD8+ CD3+ CD27− CD94+ population that induces the highest ΔY(**Fig. 5C**). We manually identified the population based on the rules specified by the decision tree model (**Fig. 5D**). We notice that the decision tree bi-sects the markers into positive and negative regions in a way that is consistent with manual gating. We previously have developed a computational tool named MetaCyto that can identify cell subsets based on their definitions^9^. We used MetaCyto to identify the CD8+ CD3+ CD27− CD94+ population across all 9 studies and found that the population is consistently increased in all studies (**Fig. 5E, Methods**).

Because of the redundancy between cell markers, cell populations can often be identified using different cell marker combinations. We further inspected the cell subset to see if the CD8+ CD3+ CD27− CD94+ population corresponds to a previously described cell population. We found that the CD8+ CD3+ CD27− CD94+ population does not correspond to any of the four well-characterized subsets of CD8+ T cells (Naive, Effector, Central Memory, and Effector Memory CD8+ T cells). Rather, all four subsets are present in the CD8+ CD3+ CD27− CD94+ population (**Fig. 5D**). Among the four subsets, the Effector and Effector memory cells are enriched in CD8+ CD3+ CD27− CD94+ cells compared to the bulk CD8+ T cells population. We then quantified the proportion of CD27− CD94+ cell subsets within the Naive, Effector, Central Memory, and Effector Memory CD8+ T cells. We found that CD27− CD94+ cells are increased in all four T cell compartments (**Fig. S4**), suggesting that CMV infection induces the CD94+ CD27− phenotype through a mechanism that is distinct from T cell activation and memory.

## Discussion

A key advantage of deep learning has been its ability to jointly optimize the feature extraction and classification steps to maximize the prediction accuracy, leading to its success in tasks involving unstructured data, such as image recognition and natural language processing^27,28^. This advantage makes the deep learning model a natural choice for analyzing cytometry data. The traditional cell-gating methods can be viewed as a way to extract features from the cytometry data. Because the cell gating step is disconnected from the later classification process, the cell gating results are often not optimized for identifying cell populations that are most associated with the outcome of interest. In the deep CNN model, the back-propagation algorithm iteratively updates the convolution layers based on classification accuracy, therefore achieving higher sensitivity in detecting cell subsets that are associated with the output of interest.

A previous study described a novel method called cellCNN^15^, which uses a single layer convolutional neural network to analyze cytometry data. While this work was innovative, a limitation of the cellCNN model is that the single convolutional layer is only able to extract cellular features by combining cell markers linearly. We extend the cellCNN model by introducing multiple convolution layers and dense layers, allowing the extraction of cellular features using complex non-linear combinations of markers. Our results show that the deep CNN model is able to identify cell populations that require multi-level hierarchical gatings, such as plasmablast, effector memory CD8+ T cells and NKT cells. In addition, the multiple layers of the deep CNN model are able to mitigate the batch effects, making the model more generalizable across studies.

In order to interpret the convolutional layers, we grouped the cells into previously defined cell subsets and quantified the mean activation value in each cell subset. We identified multiple cell subsets associated with CMV infection, including effector CD8+ T cells, Plasmablasts and CD8+ effector memory cells. Interestingly, not all the cell subsets identified from the first convolution layer are associated with CMV infection. In contrast, all the subsets identified from the second convolutional layer are significantly associated with CMV infection. The results suggest that the first convolution layer captures intermediate cellular features that do not directly correlate with CMV infection but are essential for identifying CMV-associated cell subsets in the later convolution layers.

The current study has several limitations. First, our analysis of the CMV datasets is a retrospective study. Future studies are needed to prospectively validate the diagnostic model and test the causal relationship between immune cells and CMV infection. Second, the deep neural network requires a large dataset for training, limiting its use in small scale studies. The limitation can be potentially solved by transfer learning^29^. Publicly available cytometry data can be used to pre-train the network for extracting cellular features from the markers. The last dense layers of the pre-trained model can then be trained using task-specific data for predicting the outcome of interest. Third, the current CNN model predicts the clinical outcome using cytometry data from a single time point. In many cases, the histories of the immune states are important for diagnosis or prediction. For example, the change of the immune system before and after vaccination is predictive of the vaccine responses^2,30^. In future studies, we will combine the CNN model with recurrent neural networks (RNN), such as a Long Short Term Memory (LSTM) model, to model the change of the immune system over time.

Latent infection with CMV is asymptomatic and induces limited perturbation of the immune system, making it a challenging task to diagnose the latent CMV using CyTOF data of peripheral blood samples. Despite the subtlety of changes in the immune system, the deep CNN model is able to diagnose the latent CMV infection with high accuracy. The result suggests that the deep CNN model can potentially be used to diagnose more severe conditions, including autoimmune diseases, cancer, and symptomatic infections. We envision the use of CyTOF and deep CNN as a screening tool for diagnosing a wide range of conditions, whose results can be further confirmed by established disease-specific lab tests, such as the serological test for diagnosing CMV infection^31^.

## Methods

### Data preparation

We first queried the ImmPort database to identify samples from healthy individuals with both CyTOF and CMV antibody titer data. The query identified 472 samples from nine studies, including SDY112, SDY113, SDY305, SDY311, SDY472, SDY472, SDY515, and SDY519 as of March 2019^16–19^. We downloaded CyTOF data and transformed the raw cytometry signal using arcsinh transformation (y = arcsinh(x/5)). To combine CyTOF samples, we included 28 markers that are present in data from all nine studies and subsampled 10000 cells from each sample. The final CyTOF data are organized into a three-dimensional matrix (472 samples × 28 markers × 10000 cells).

### Deep CNN architecture

The deep CNN model takes cytometry matrices as inputs. For each sample, the matrix profiles multiple markers (columns) for single cells (rows). Convolution layers are used after the input layer to extract cellular features from the cytometry data. The filter size in the first convolution layer is 1 × m × 1, where m is the number of markers in cytometry data. The filter size used in the subsequent convolutional layers is 1 × 1 × f, where f is the number of filters in the previous convolution layer. The cellular features of the last convolution layer are pooled into sample level features using either max or mean pooling. The pooling layer is followed by dense layers, which combine the features extracted by the convolutional layers and summarized by the pooling layers. In the output layer, a logistic regression combines the output of the last dense layer to predict binary outcomes. For continuous outcomes, linear regression is used. For each layer, batch normalization is used for regularization and to facilitate model training. We used Rectified Linear Unit (ReLU) as the activation function for all internal layers.

### Training, optimization, and testing of the deep CNN model

We used the Adam algorithm, a variant of the gradient descent, to identify the best parameters in the neural network^32^, with binary cross-entropy as the loss function. To prevent overfitting, the performance of the model is tested at each epoch using the validation data. The parameters that give rise to the best validation result are used in the final model.

The hyperparameters of the deep learning model include the number of convolution layers, the number of filters in the convolution layers, the type of pooling layer (max or mean pooling), the number and size of the dense layers and the learning rate. We performed a grid search to optimize hyperparameters using the training and validation datasets. The optimized model for diagnosing CMV contains two convolution layers with three filters in each layer, a mean pooling layer, and a three-node dense layer. The model is trained with a learning rate of 0.0001, batch size of 60 and total epochs of 500. The performance of the optimized model is tested using the test dataset (SDY519), which has not been used during the training and optimization processes.

### Training and optimization of CytoDx, CellCNN, and FlowSOM

To test the performance of CytoDx, CellCNN and FlowSOM, we used the same training, validation and testing datasets that had been applied to the deep CNN model (**Fig. 2A**). We trained two CytoDx models using the CytoDx R package. The first model uses the arcsinh transformed cytometry data as input. The second model uses the rank-transformed cytometry data and the two-way interactions between each pair of markers. We used the validation dataset to evaluate the two models and found the second model to be superior. We benchmarked its performance using the test dataset.

We performed a grid search for the CellCNN model (number of filters ranging from 2-10, drop out rate ranging from 0.1 to 0.9). Using the validation dataset, we chose an optimal set of hyper-parameters (number of filters equals 5, drop out rate equals 0.2). Adam algorithm is used for training the model with a learning rate of 0.001. The trained model is evaluated using the testing dataset.

Using FlowSOM, we clustered the cells data using a 10-by-10 self-organizing map (SOM) and identified 20 meta-clusters from the SOM result. We derived summary statistics of the identified cell subsets, including percentage in PBMC and mean fluorescence intensity (MFI) of cell markers. We then trained a Random Forest model (number of trees = 100) to predict the latent CMV infection in the subjects using results from FlowSOM as input. The optimized models were evaluated using the testing dataset.

### Measurement of the heterogeneity between datasets

We calculated the average marker intensities of each sample as a surrogate to measure heterogeneity between studies. For internal layers of the deep CNN model, we calculated the average activation value of each sample in each layer. We then use the Kruskal–Wallis test (also known as the one-way ANOVA on ranks) to test if the average marker or activation values are significantly different between studies. We used the non-parametric Kruskal-Wallis test because the activation values are not normally distributed due to the use of Relu and logistic activation functions.

### Quantifying activation value in cell populations

We extracted the activation values of the internal nodes in each filter of the convolutional layers. Using definitions from the Human Immunology Project Consortium, We identified 24 immune cell subsets from the CyTOF data. We mapped the activation values to the 24 cell populations and calculated the mean activation value for each population. We normalized the mean activation value to the maximum activation value in each convolutional layer.

### Permutation based interpretation of deep CNN model

For each cell in cytometry data, we up-sampled the cell by copying it to replace other randomly chosen cells within the sample. We then applied the deep CNN model on both the original data and the permuted data. The difference in the model output (ΔΥ) quantifies the impact of the cell on the output of the deep learning model. We repeated the process until ΔΥ is calculated for all cells in SDY519.

We choose to up-sample the cells, rather than delete the cell, to evaluate its impact. This is because cytometry data contains a large number of cells, deleting a single cell has an extremely limited impact on the model output. On the other hand, we can up-sample the cell to replace a significant proportion of cells in the sample, therefore inducing a significant change to the model output. We up-sampled every cell to 1% or 5% of the total population and found that the ΔY are highly correlated between the two scenarios, suggesting that the ΔY is robust to the level of upsampling (**Fig. S2**). We chose to up-sample each cell to 5% of the sample in this study.

Decision trees were trained using the CyTOF data as inputs, and ΔY as outputs. The DecisionTreeClassifier function in the scikit-learn package is used to construct the decision tree. To determine the depth of the decision tree, we constructed decision trees with maximum depth from 2 to 10. We measured the performance of the decision trees using the correlation between observed ΔY and fitted ΔY. We used the “elbow” method and determined an optimal depth of 4 (**Fig. S3**).

### Quantify cell populations using MetaCyto

To identify the cell subset with the highest ΔY, we inspected the decision tree model and identified the hierarchical decision rule that leads to the leaf with the highest mean ΔY (CD8>2.4, CD27<1.38, CD3>2.14, CD94>.82). We notice that the decision tree bi-sect the markers into positive and negative regions in a way that is consistent with manual gating. We, therefore, specified the cell definition to be CD8+ CD27− CD3+ CD94+, which can be used as input in our previously developed MetaCyto R package. Using the “searchCluster” function in MetaCyto, we quantified the proportion of CD8+ CD27− CD3+ CD94+ subset across nine studies. Using the same procedure, we quantified the proportion of CD27− CD94+ cells within Naive, effector, effector memory and central memory CD8+ T cells.

### Statistical analysis

We performed 1000-fold bootstrapping to test if the performances of two machine learning models are equal. In each iteration, we sampled from the testing dataset with replacement and evaluated the performance of the two models using AUC. We calculated the p-value as the percentage of interactions in which a model outperforms the other. We measured the batch effect in each layer using the Kruskal–Wallis test. We used a two-way ANOVA model to test the association between a cell subset and CMV infection, in which the proportion of the cell subset is regressed on CMV infection and study.

### Availability of Data and code

The CyTOF and anti-CMV antibody titer data are publically available on ImmPort. We provided a tutorial demonstrating how to create, train and interpret the deep CNN model (https://github.com/hzc363/DeepLearningCyTOF). All the codes used in the study are available on GitHub (https://github.com/hzc363/Deep_learning_CyTOF_Code).

## Acknowledgments

We would like to thank Mark Davis and David Hafler for openly sharing their CyTOF measurement data on the ImmPort database, facilitating research like ours. We would like to thank Patrick Dunn, Elizabeth Thomson, Henry Schaefer, Daniel Wong, Dmytro Lituiev, Benjamin Glicksberg and Matthew Elliott for helpful discussions. We thank Boris Oskotsky for server support. The research reported in this publication was supported by the National Institute of Allergy and Infectious Diseases (Bioinformatics Support Contract HHSN272201200028C). The content is solely the responsibility of the authors and does not necessarily represent the official views of the National Institutes of Health.

## Author contributions

Z.H. designed and implemented the deep learning model. Z. H., A. T., and J. S. performed the analysis to test and interpret the deep learning model. S. B. and A. B. gave valuable input and suggestions for analysis. Z.H and A. B. led the study. All authors wrote the manuscript.

## Figure Legends

**Supplementary Figure 1:** Box Plots showing the percentage of six cell populations in CMV+ and CMV-subjects across nine studies. The cell subsets are selected based on their high activation value in convolutional layers (See **Fig. 5A**). P values are from two-way ANOVA models, with CMV infection and study as two factors. The p values of the CMV infection variable are reported.

**Supplementary Figure 2:** The change of output from the deep CNN model (ΔY) is measured by up-sampling every single cell in the test dataset to 1% or 5% of the total population. The scatter plot shows the correlation between the two sets of ΔY.

**Supplementary Figure 3:** The performance of the decision trees are measured by the correlation between the input data (ΔY of the deep CNN model, see **Fig. 5B**) and output. The plot shows the relationship between the correlation and the maximum depth of the decision trees.

**Supplementary Figure 4:** Proportion of CD27− CD94+ subset within CD8+ Naive, Effector, Central Memory and Effect Memory T cell compartments. P values are from two-way ANOVA models, with CMV infection and study as two factors. The p values of the CMV infection variable are reported.

**Supplementary Table 1:** The performance of the deep CNN model in 10-interaction permutation analysis. In each iteration, we randomly assigned one study as the validation dataset, one study as the test dataset and the rest studies as the training dataset to train and evaluate the deep CNN model.

## References

1. Spitzer, M. H. & Nolan, G. P. Mass Cytometry: Single Cells, Many Features. Cell 165, 780–791 (2016).

2. Nakaya, H. I. et al. Systems biology of vaccination for seasonal influenza in humans. Nat. Immunol. 12, 786–795 (2011).

3. Michlmayr, D. et al. Comprehensive innate immune profiling of chikungunya virus infection in pediatric cases. Mol. Syst. Biol. 14, e7862 (2018).

4. Spitzer, M. H. et al. Systemic Immunity Is Required for Effective Cancer Immunotherapy. Cell 168, 487–502.e15 (2017).

5. Rawstron, A. C. et al. Reproducible diagnosis of chronic lymphocytic leukemia by flow cytometry: An European Research Initiative on CLL (ERIC) & European Society for Clinical Cell Analysis (ESCCA) Harmonisation project. Cytometry B Clin. Cytom. 94, 121–128 (2018).

6. Ocmant, A. et al. Flow cytometry for basophil activation markers: The measurement of CD203c up-regulation is as reliable as CD63 expression in the diagnosis of cat allergy. J. Immunol. Methods 320, 40–48 (2007).

7. Farias, M. G., de Lucena, N. P., Dal Bó, S. & de Castro, S. M. Neutrophil CD64 expression as an important diagnostic marker of infection and sepsis in hospital patients. J. Immunol. Methods 414, 65–68 (2014).

8. Qian, Y. et al. Elucidation of seventeen human peripheral blood B-cell subsets and quantification of the tetanus response using a density-based method for the automated identification of cell populations in multidimensional flow cytometry data. Cytometry B Clin. Cytom. 78B, S69–S82 (2010).

9. Hu, Z. et al. MetaCyto: A Tool for Automated Meta-analysis of Mass and Flow Cytometry Data. Cell Rep. 24, 1377–1388 (2018).

10. Van Gassen, S. et al. FlowSOM: Using self-organizing maps for visualization and interpretation of cytometry data. Cytom. Part J. Int. Soc. Anal. Cytol. 87, 636–645 (2015).

11. Bruggner, R. V., Bodenmiller, B., Dill, D. L., Tibshirani, R. J. & Nolan, G. P. Automated identification of stratifying signatures in cellular subpopulations. Proc. Natl. Acad. Sci. U. S. A. 111, E2770–E2777 (2014).

12. Gassen, S. V., Vens, C., Dhaene, T., Lambrecht, B. N. & Saeys, Y. FloReMi: Flow density survival regression using minimal feature redundancy. Cytometry A 89, 22–29 (2016).

13. Choileain, S. N. et al. The Dynamic Processing of CD46 Intracellular Domains Provides a Molecular Rheostat for T Cell Activation. PLOS ONE 6, e16287 (2011).

14. Hu, Z., Glicksberg, B. S. & Butte, A. J. Robust prediction of clinical outcomes using cytometry data. Bioinformatics 35, 1197–1203 (2019).

15. Arvaniti, E. & Claassen, M. Sensitive detection of rare disease-associated cell subsets via representation learning. Nat. Commun. 8, 14825 (2017).

16. Bhattacharya, S. et al. ImmPort, toward repurposing of open access immunological assay data for translational and clinical research. Sci. Data 5, 180015 (2018).

17. Kronstad, L. M., Seiler, C., Vergara, R., Holmes, S. P. & Blish, C. A. Differential Induction of IFN-α and Modulation of CD112 and CD54 Expression Govern the Magnitude of NK Cell IFN-γ Response to Influenza A Viruses. J. Immunol. 201, 2117–2131 (2018).

18. Miron, M. et al. Human Lymph Nodes Maintain TCF-1hi Memory T Cells with High Functional Potential and Clonal Diversity throughout Life. J. Immunol. 201, 2132–2140 (2018).

19. Alpert, A. et al. A clinically meaningful metric of immune age derived from high-dimensional longitudinal monitoring. Nat. Med. 25, 487 (2019).

20. Rusu, R. B., Marton, Z. C., Blodow, N., Dolha, M. & Beetz, M. Towards 3D Point cloud based object maps for household environments. Robot. Auton. Syst. 56, 927–941 (2008).

21. Qi, C. R., Su, H., Mo, K. & Guibas, L. J. PointNet: Deep Learning on Point Sets for 3D Classification and Segmentation. ArXiv161200593 Cs (2016).

22. van Boven, M. et al. Infectious reactivation of cytomegalovirus explaining age- and sex-specific patterns of seroprevalence. PLoS Comput. Biol. 13, (2017).

23. Colugnati, F. A., Staras, S. A., Dollard, S. C. & Cannon, M. J. Incidence of cytomegalovirus infection among the general population and pregnant women in the United States. BMC Infect. Dis. 7, 71 (2007).

24. Fowler, K. B. et al. Racial and Ethnic Differences in the Prevalence of Congenital Cytomegalovirus Infection. J. Pediatr. 200, 196–201.e1 (2018).

25. Finak, G. et al. Standardizing Flow Cytometry Immunophenotyping Analysis from the Human ImmunoPhenotyping Consortium. Sci. Rep. 6, 20686 (2016).

26. Ribeiro, M. T., Singh, S. & Guestrin, C. ‘Why Should I Trust You?’: Explaining the Predictions of Any Classifier. (2016).

27. Lai, S., Xu, L., Liu, K. & Zhao, J. Recurrent Convolutional Neural Networks for Text Classification. in Twenty-Ninth AAAI Conference on Artificial Intelligence (2015).

28. He, K., Zhang, X., Ren, S. & Sun, J. Deep Residual Learning for Image Recognition. in 2016 IEEE Conference on Computer Vision and Pattern Recognition (CVPR) 770–778 (2016). doi:10.1109/CVPR.2016.90.

29. Shin, H. et al. Deep Convolutional Neural Networks for Computer-Aided Detection: CNN Architectures, Dataset Characteristics and Transfer Learning. IEEE Trans. Med. Imaging 35, 1285–1298 (2016).

30. Vahey, M. T. et al. Expression of genes associated with immunoproteasome processing of major histocompatibility complex peptides is indicative of protection with adjuvanted RTS,S malaria vaccine. J. Infect. Dis. 201, 580–589 (2010).

31. Ross, S. A., Novak, Z., Pati, S. & Boppana, S. B. Diagnosis of Cytomegalovirus Infections. Infect. Disord. Drug Targets 11, 466–474 (2011).

32. Kingma, D. P. & Ba, J. Adam: A Method for Stochastic Optimization. ArXiv14126980 Cs (2014).

