## Supplementary material for "A robust and interpretable, end-to-end deep learning model for cytometry data": Fig. S1

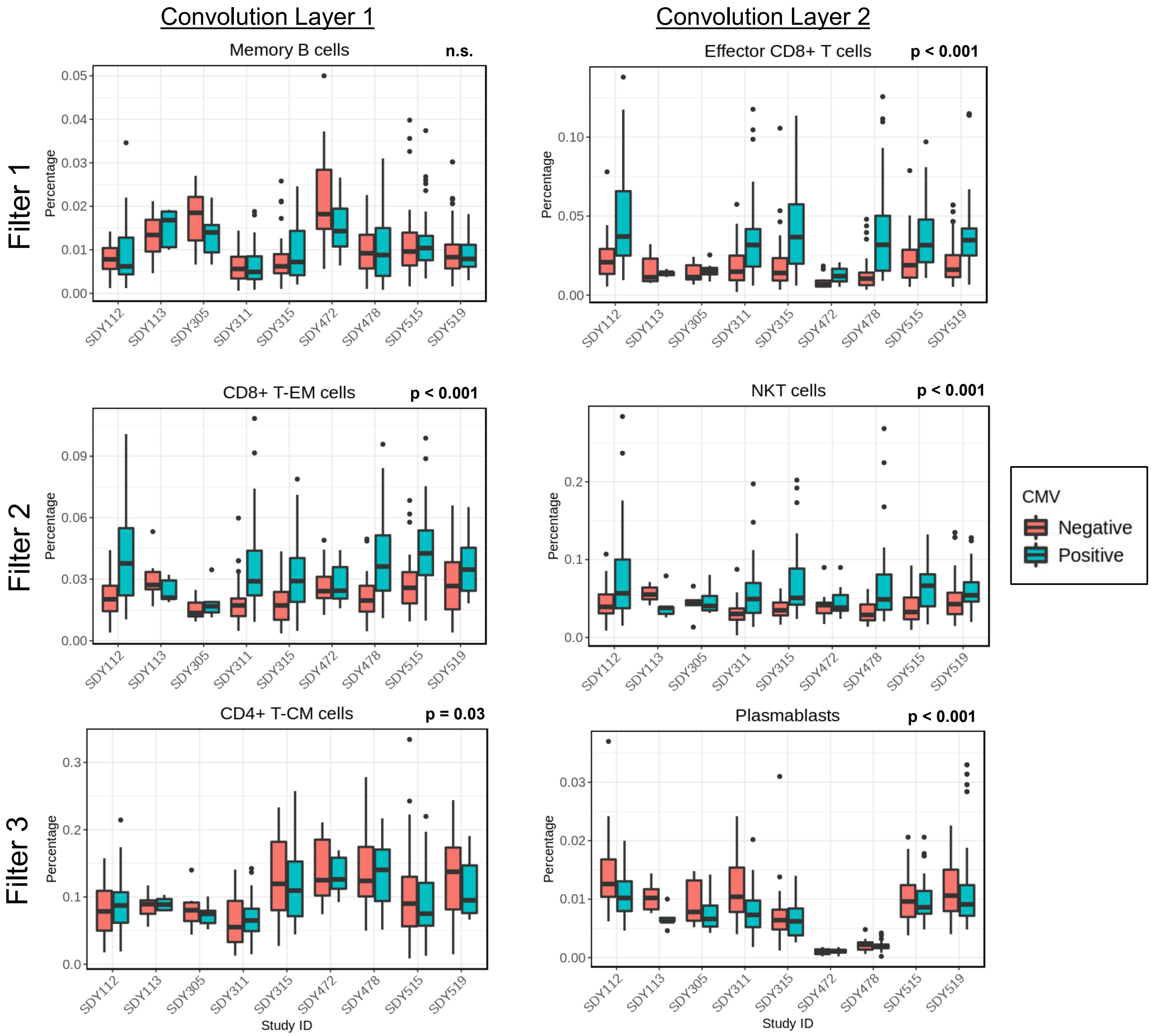


**Supplementary Figure 1:** Box Plots showing the percentage of six cell populations in CMV+ and CMV- subjects across nine studies. The cell subsets are selected based on their high activation value in convolutional layers (**See Fig. 5A**). P values are from two-way ANOVA models, with CMV infection and study as two factors. The p values of the CMV infection variable are reported.
