## Supplementary material for "A robust and interpretable, end-to-end deep learning model for cytometry data": Fig. S2

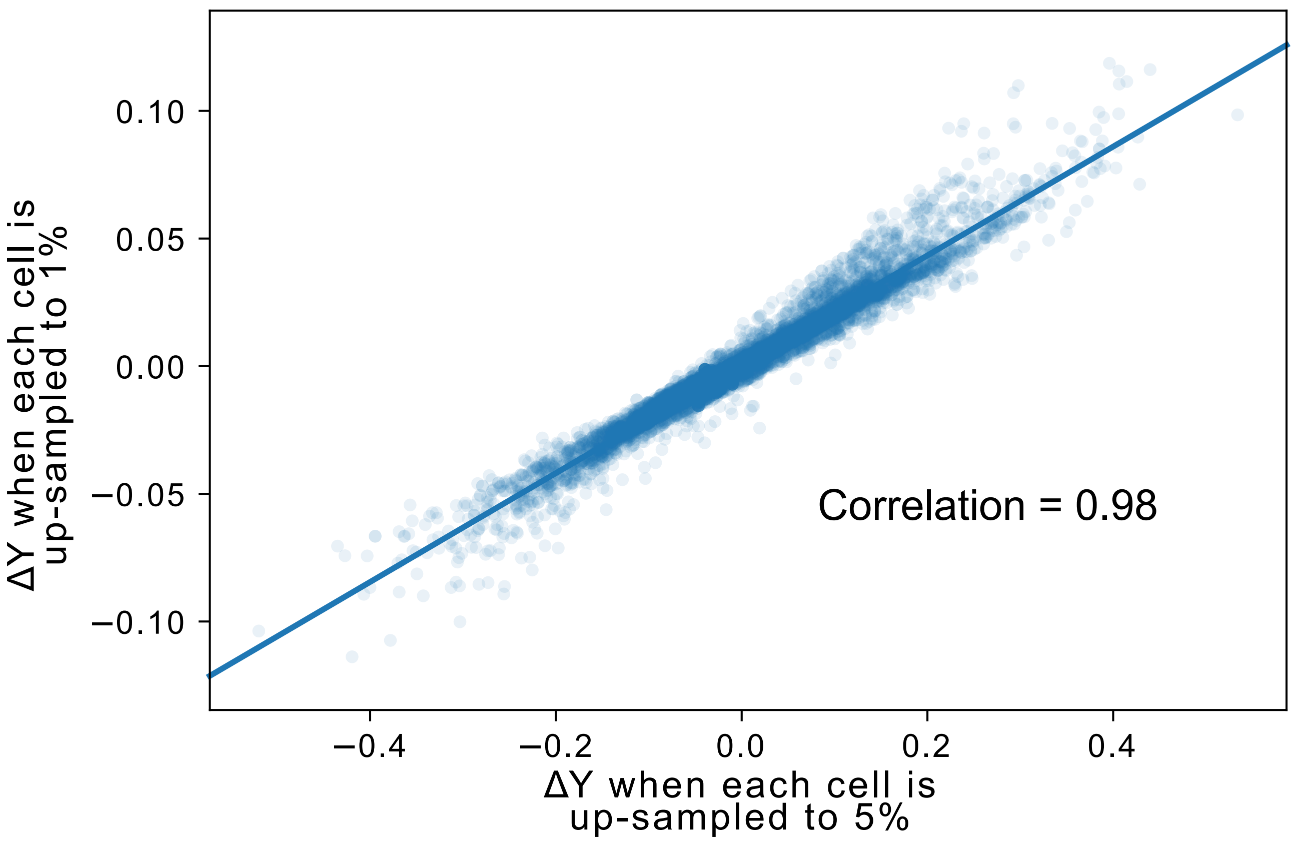


**Supplementary Figure 2:** The change of output from the deep CNN model (ΔY) is measured by up-sampling each single cell in the test dataset to 1% or 5% of the total population. The scatter plot shows the correlation between the two sets of ΔY.
