## Supplementary material for "A robust and interpretable, end-to-end deep learning model for cytometry data": Fig. S3

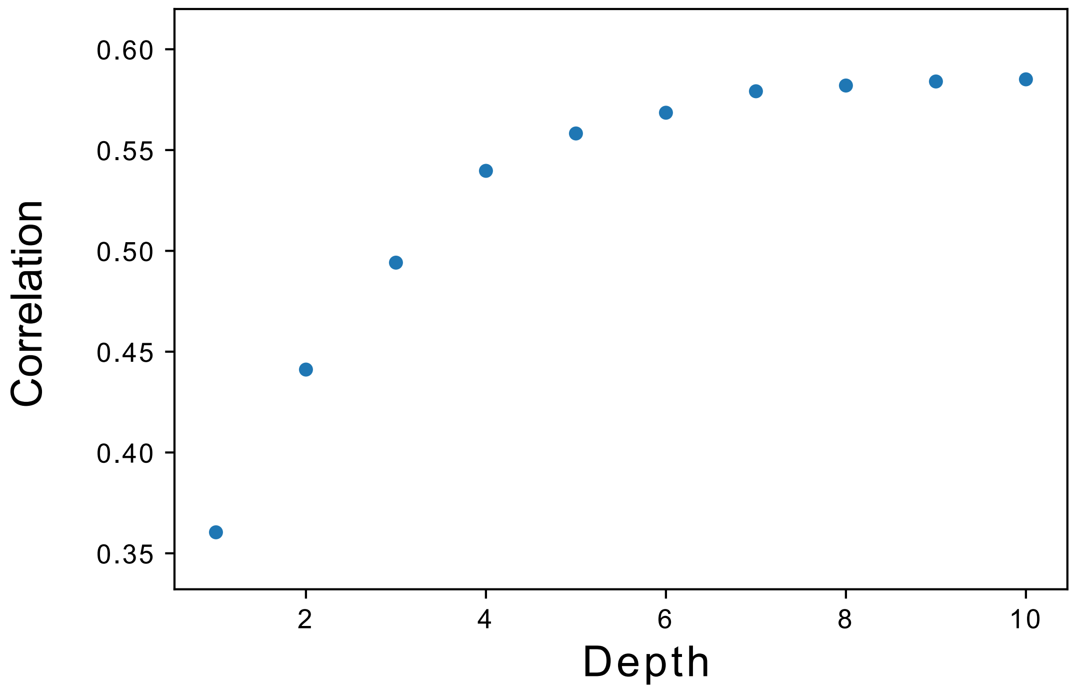


**Supplementary Figure 3:** The performance of the decision trees are measured by the correlation between the input data (ΔY of the deep CNN model, see **Fig. 5B**) and output. The plot shows the relationship between the correlation and the maximum depth of the decision trees.
