## Supplementary material for "A robust and interpretable, end-to-end deep learning model for cytometry data": Fig. S4

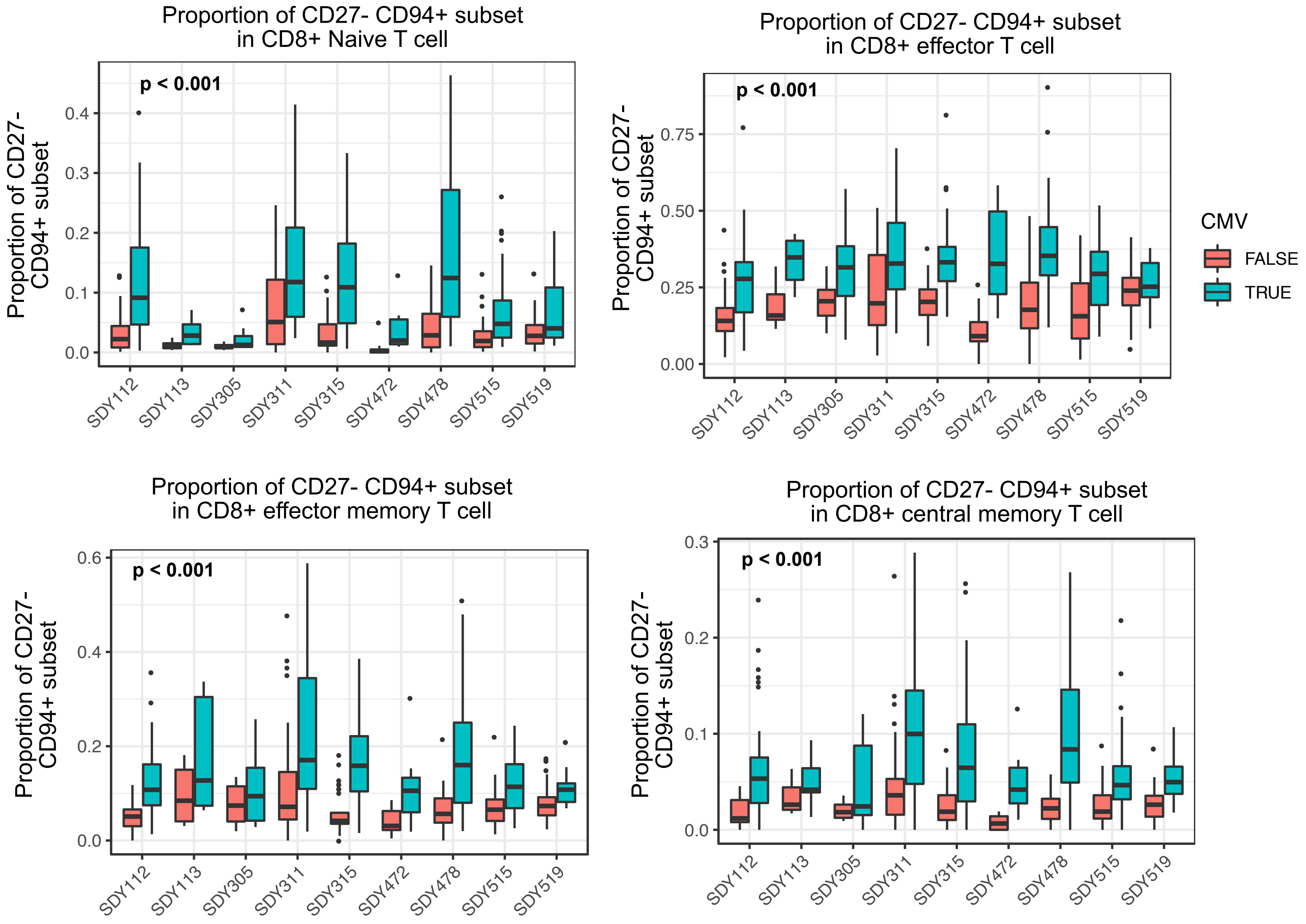


**Figure S4:** Proportion of CD27- CD94+ subset within CD8+ Naive, Effector, Central Memory and Effect Memory T cell compartment. P values are from two-way ANOVA models, with CMV infection and study as two factors. The p values of the CMV infection variable are reported.
