## Supplementary material for "A robust and interpretable, end-to-end deep learning model for cytometry data": Table S1

| Validation | Test | AUROC |
|---|---|---|
| SDY305 | SDY113 | 0.93 |
| SDY311 | SDY478 | 0.94 |
| SDY472 | SDY478 | 0.96 |
| SDY519 | SDY112 | 0.94 |
| SDY515 | SDY519 | 0.95 |
| SDY519 | SDY113 | 0.97 |
| SDY311 | SDY315 | 0.95 |
| SDY112 | SDY315 | 0.95 |
| SDY472 | SDY519 | 0.99 |
| SDY311 | SDY515 | 0.95 |

**Supplementary Table 1:** The performance of the deep CNN model in 10-interaction permutation analysis. In each iteration, we randomly assigned one study as the validation dataset, one study as the test dataset and the rest studies as the training dataset to train and evaluate the deep CNN model.
